# Emergence and disruption of cooperativity in a denitrifying microbial community

**DOI:** 10.1101/2024.10.24.620115

**Authors:** Alex V. Carr, Anne E. Otwell, Kristopher A. Hunt, Yan Chen, James Wilson, José P. Faria, Filipe Liu, Janaka N. Edirisinghe, Jacob J. Valenzuela, Serdar Turkarslan, Lauren M. Lui, Torben N. Nielsen, Adam P. Arkin, Christopher S. Henry, Christopher J. Petzold, David A. Stahl, Nitin S. Baliga

## Abstract

Anthropogenic perturbations to the nitrogen cycle, primarily through use of synthetic fertilizers, is driving an unprecedented increase in the emission of nitrous oxide (N_2_O), a potent greenhouse gas and an ozone depleting substance, causing urgency in identifying the sources and sinks of N_2_O. Microbial denitrification is a primary contributor to biotic production of N_2_O in anoxic regions of soil, marine systems, and wastewater treatment facilities. Here, through comprehensive genome analysis, we show that pathway partitioning is a ubiquitous mechanism of complete denitrification within microbial communities. We have investigated mechanisms and consequences of process partitioning of denitrification through detailed physiological characterization and kinetic modeling of a synthetic community of *Rhodanobacter R12* and *Acidovorax 3H11*. We have discovered that these two bacterial isolates, from a heavily nitrate (NO_3_^−^) contaminated superfund site, complete denitrification through the exchange of nitrite (NO_2_^−^) and nitric oxide (NO). The process partitioning of denitrification and other processes, including amino acid metabolism, contribute to increased cooperativity within this denitrifying community. We demonstrate that certain contexts, such as high NO_3_^−^, cause unbalanced growth of community members, due to differences in their substrate utilization kinetics. The altered growth characteristics of community members drives accumulation of toxic NO_2_^−^, which disrupts denitrification causing N_2_O off gassing.

## INTRODUCTION

Since the development of the Haber-Bosch process, there has been an exponential increase in the global deposition of fixed nitrogen (N). This has largely resulted from the use of synthetic fertilizers^1^. Meeting the demands of a growing global population has contributed to a nearly ten-fold increase in synthetic fertilizer use from 1960 to 2013^2^. Recent estimates suggest anthropogenic contributions account for about half of reactive N flux on Earth^2^. As a result, the global N cycle has become increasingly perturbed, contributing to eutrophication of terrestrial and aquatic systems, global acidification, increasing atmospheric nitrous oxide (N_2_O) and stratospheric ozone loss^1^. The microbial processes of nitrification and denitrification are responsible for the bulk of N_2_O emissions in agricultural systems and wastewater treatment^3–5^. Thus, understanding how microbial processes, interactions, and environmental factors such as resource concentration, pH, and metal availability influence the fate of N compounds, and the production of N_2_O in particular, is essential for developing a predictive understanding of the fate of different N species in natural and engineered systems^6–9^.

Denitrification is the major biological process that returns fixed N back to the atmosphere through reduction of N-oxides nitrate (NO_3_^−^) and nitrite (NO_2_^−^) to gaseous compounds nitric oxide (NO), N_2_O, and dinitrogen (N_2_). Therefore, it comprises a critical step in the global N cycle. Denitrification is also an anaerobic process, which provides an alternative means of generating energy when oxygen (O_2_) is not available. In complete denitrification, microorganisms reduce NO_3_^−^ to N_2_ in a series of four reductive, energy-conserving steps. First, NO_3_^−^ is reduced to NO_2_^−^ by one of two dissimilatory nitrate reductases—the membrane-bound Nar complex or the periplasmic nitrate reductase, Nap. NO_2_^−^ is then reduced to NO by one of two structurally unrelated nitrite reductases that contain different prosthetic groups—the copper-containing NirK or the cytochrome cd1 NirS^10^. NO is then reduced to N_2_O by the nitric oxide reductase (NorB), and finally N_2_O is reduced to N_2_ by the nitrous oxide reductase (NosZ), of which two distinct clades have been identified^10,11^.

Most studies have only examined complete denitrifiers^12,13^ in monoculture to study this respiratory process (enzymology, kinetics, regulation, etc.). However, ongoing genomic and metagenomic characterization of isolates and natural communities has revealed, remarkably, that the genomes of most organisms that encode denitrification enzymes lack the complete pathway^14^. More commonly, the microbial communities in soil that drive denitrification are composed of a complex mixture of species that individually encode partial (rather than complete) denitrification pathways^15–18^. The environmental significance of organismal patchiness of genes in the pathway for denitrification is mostly unexplored but may be associated with increased efficiency and resiliency of denitrifying communities. Recent work has shown that pathway partitioning can decrease inter-enzyme competition within individual microbes and reduce the accumulation of toxic intermediates like NO_2_^−^ through their exchange, thus providing a potential selection mechanism for pathway partitioning in denitrifying communities^19^. However, understanding how this generalizes when multiple soluble N intermediates (i.e., NO_2_^−^, NO, N_2_O) are exchanged as well as how the environment selects for natural communities with specific partial pathway combinations remains unexplored. Additionally, because many soil and groundwater dwelling organisms are facultative anaerobes, the ability of these organisms to leverage multiple pathways and substrates for respiration may also contribute to denitrification pathway patchiness.

Here, we have investigated the role of pathway partitioning in a synthetic community (SynCom) composed of two isolates recovered from the same field site location at the Oak Ridge Reservation (ORR) (https://enigma.lbl.gov)^20^ and representative of naturally occurring populations capable of exchange of pathway intermediates. This two-organism SynCom served as a model for examining the roles of biotic and abiotic factors controlling N_2_O emissions in the low-oxygen and anaerobic zones of the subsurface environment of the ORR, which have high concentrations of NO_3_^−^ due to past nuclear waste disposal and evidence of high levels of denitrification activity^20,21^. Using this SynCom as a model system, we have discovered how process partitioning of denitrification and exchange of multiple N intermediates improved community growth characteristics via changes in pathway kinetics (e.g., increased rate of NO_3_^−^ reduction, removal of NO_2_^−^, and exchange of N_2_O). We also show the importance of community composition in determining the physiology and growth dynamics of each species and how changes in environmental conditions can break the collaboration between the two members, leading to N_2_O emissions.

## RESULTS

### Denitrification pathway is partitioned across members of microbial communities at the ORR

We sought to understand the structure and activity of communities responsible for denitrification at the ORR and their relation to N_2_O production. We first investigated the distribution of denitrification pathway enzymes within the genomes of 252 isolates obtained from groundwater wells of various depths distributed across 3 major areas of the ORR ^20–23^. The sites spanned areas of high, medium, and low average groundwater NO_3_^−^ concentration (between 0-250 mM NO_3_^−^). For the pathway analysis we considered the genes Nar (or Nap), Nir, Nor, and Nos. Collectively, the reactions encoded by these genes convert NO_3_^−^ to N_2_ (Fig. 1A). Genomic analysis revealed that the majority (∼71%) of isolates were incomplete denitrifiers (i.e., missing one or more genes of the pathway). Of these, 40% likely contributed to N_2_O production due to the presence of one or more genes encoding Nor but none encoding Nos. These conclusions were consistent with the pathway compositions of 4622 IMG soil isolates, a vast majority of which (90%) were also incomplete denitrifiers (Fig. 1A). Thus, the genomic analysis demonstrated that the denitrification pathway is largely partitioned across soil and ground water dwelling organisms and suggested that complete denitrification likely occurs through interactions among organisms with complementary capabilities within microbial communities.

**Figure 1.**
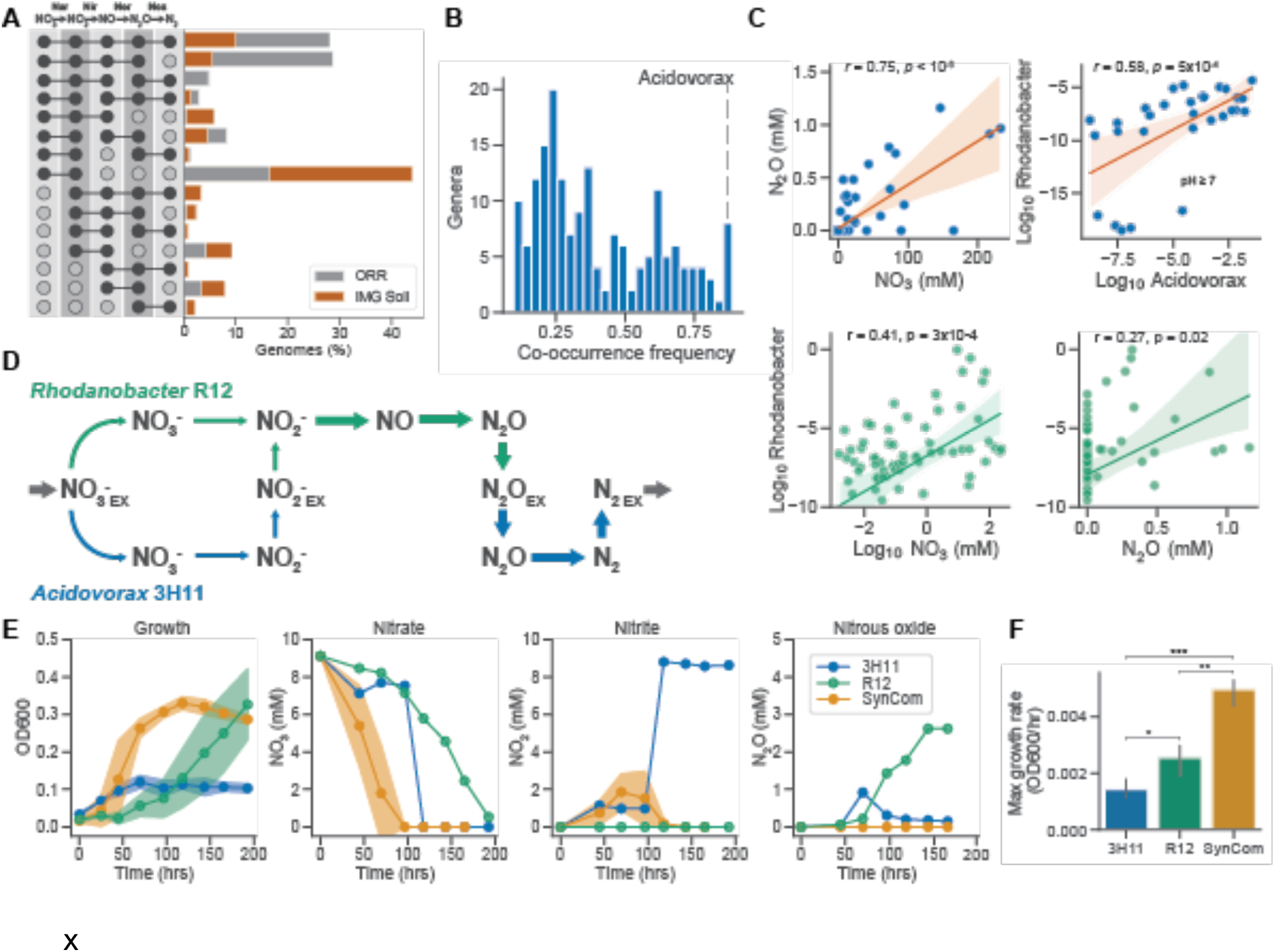
Field observations and metabolic modeling predict exchange of intermediates in a denitrifying synthetic community. (**A**) Distribution of denitrification pathway compositions among ORR field and IMG soil isolates. (**B**) Genus level co-occurrence frequency of *Rhodanobacter* and other genera across ORR groundwater samples. (**C**) Association between groundwater chemistries and genus abundance. Pearson correlation *r* and *p* values are listed in each subplot. Linear fits to data using OLS are displayed as well as 95% confidence intervals (**D**) Denitrification flux patterns predicted by genome scale metabolic model of R12 and 3H11 SynCom assuming equal abundances. (**E**) Growth characteristics of R12, 3H11 and R12-3H11 SynCom. Shading around trend lines represents standard deviation of samples and points represent averages. (**F**) Comparison of monoculture and SynCom maximum growth rates. Bars indicate comparisons for which differences were significant using Welch’s t-test. *, *p*<0.05; **, *p*<0.01; ***, *p*<0.001.

To better understand the role of pathway partitioning in denitrifying communities, we assembled the field isolates *Rhodanobacter sp.* FW510-R12 (R12) and *Acidovorax sp.* GW101-3H11 (3H11) into a SynCom capable of complete denitrification through pathway intermediate exchange. The selection of organisms in this pairing was based on relationships among genus abundance, groundwater chemistry (*e.g.*, NO_3_^−^ and N_2_O concentrations), and isolate co-occurrence determined through genomic and 16S rRNA profiling across the field site (Fig. 1B and C). We observed significant correlation between NO_3_^−^ and N_2_O concentrations in the ORR groundwater (Fig. 1C), consistent with previous observations that high NO_3_^−^ can lead to N_2_O emissions^3,20,22^. We also found that the abundance of *Rhodanobacter spp*. was correlated with both NO_3_^−^ and N_2_O levels at the field site (Fig. 1C), which was consistent with the reported high abundance of this genus in regions of high NO_3_^−^ and heavy metal contamination in the ORR^8,23^. While the abundance of *Acidovorax spp.* was weakly associated with NO_3_^−^ and N_2_O concentrations (Fig. 1C), it was positively correlated with *Rhodanobacter spp*. (Fig. 1C) in non-acidic groundwater (pH≥7). Flux balance analysis using constraints-based metabolic models indicated that a SynCom of R12 and 3H11 could perform complete denitrification through the exchange of NO_2_^−^ and N_2_O (Fig. 1D). Consistent with model predictions, 3H11 and R12 were both capable of growth on anaerobic minimal medium containing NO_3_^−^ and acetate as the primary electron acceptor and donor pair, respectively. Additionally, consistent with the absence of a genome-encoded NO_2_^−^ reductase (Nir), 3H11 consumed NO_3_^−^ and accumulated NO_2_^−^ (Fig. 1E). By contrast, R12 consumed NO_3_^−^, and accumulated N_2_O with no transient accumulation of NO_2_^−^, implying that the rate of NO_2_^−^ reduction was equal to or greater than the rate of NO_3_^−^ reduction (Fig. 1E).

The SynCom growth rate was significantly greater than that of monocultures of either of the two isolates (Fig. 1F). Additionally, the rate of NO_3_^−^ reduction was increased relative to the individual monocultures, followed by a transient accumulation of NO_2_^−^, which was fully reduced to N_2_ gas based on the lack of N_2_O and ammonia accumulation (Fig. 1E). In depth growth characterization demonstrated that 3H11 and R12 complemented each other to perform complete denitrification, which manifested in synergistic improvement in overall growth dynamics relative to monocultures of the two organisms.

### Exchange of intermediates in the denitrification pathway largely explain enhanced growth phenotype of the SynCom

We hypothesized that synergistic improvement in rate of NO_3_^−^ reduction and overall growth characteristics of the SynCom were likely due to R12-mediated rescue of NO_2_^−^-mediated growth inhibition of 3H11, which in turn reduced N_2_O to N_2_. To test this hypothesis, we characterized the growth kinetics of each monoculture in media supplemented with varied concentrations of NO_3_^−^, NO_2_^−^, and N_2_O (Fig. S1) and used that data to develop kinetic models of growth dynamics of each isolate and the SynCom. The growth rate of 3H11 was significantly greater than R12 across a wide range of NO_3_^−^ (Fig. 2A), achieving its maximum growth rate at a much lower concentration (Ks_3H11_<<Ks_R12_, Welch’s t-test *t*=26.1, *p<*10^−6^, Fig. S1F). While a low concentration of NO_2_^−^ supported a significantly higher growth rate of R12 relative to its growth rate on NO_3_^−^ (10-fold greater for 1-5 mM substrate, Fig. 2A), growth was reduced at higher concentrations (Ki_R12_=13.23 mM, Fig. S1F) and completely inhibited at 10 mM NO_2_^−^ (Fig. S1B and D). Nitrite also had a strong inhibitory effect on the growth rate and biomass yield of 3H11, with complete inhibition above 5 mM NO_2_^−^ in media containing 10 mM NO_3_^−^ (Ki_3H11=_9.11 mM; Figs. 2A and S1B, D and F). Notably, R12 appeared to not conserve energy from the reduction of NO_3_^−^, since yield and growth rates were identical on NO_3_^−^ and NO_2_^−^ (slope_NO3_=0.033 ± 0.003, slope_NO2_=0.032 ± 0.006 for ordinary least squares fits to linear portion of substrate versus maximum OD600; substrate concentrations 1-10 mM; Welch’s t-test *t*=1.1, *p=*0.27). Thus, R12 growth was modeled as entirely dependent on the reduction of NO_2_^−^ and NO. Finally, while 3H11 was able to accumulate biomass by reducing N_2_O, its growth rate on N_2_O was constant and not dose-dependent (2-30 mM; Ks_3H11_ ≈ 0, Fig. 2A and S1D and F).

**Figure 2.**
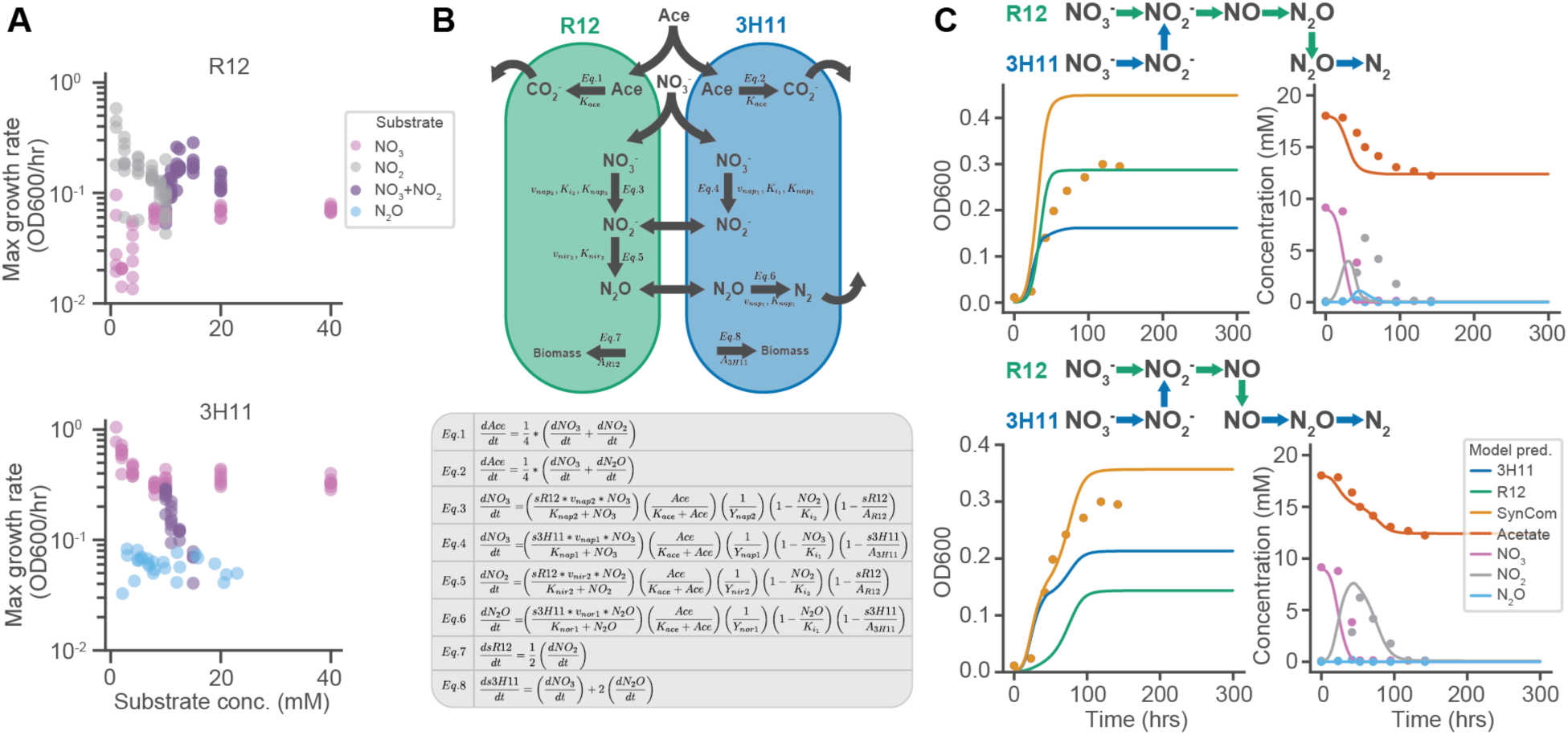
Exchange of pathway intermediates drives growth enhancement in R12-3H11 SynCom. (**A**) Maximum growth rates of R12 and 3H11 across variation in substrate concentrations. For conditions where NO_3_^−^ and NO_2_^−^ were present, 10 mM NO_3_^−^ was used and NO_2_^−^ was varied. (**B**) Schematic of R12-3H11 kinetic model and associated differential equations. Model represents growth resulting from oxidation of acetate and reduction of NO_3_^−^, NO_2_^−^, and N_2_O via denitrification. Given challenges of measuring NO kinetics, NO reduction is not explicitly represented in the model, instead NO_2_^−^ is reduced to N_2_O (**C**) Two SynCom intermediate exchange scenarios and associated dynamics predicted by kinetic models displayed with empirical data. Model predictions are represented using solid lines, data are averages across samples and are represented using circular points.

Kinetic models were developed based on a modified Monod framework, integrating logistic representation of carrying capacity into equations describing growth kinetics as a function of metabolite concentrations (Fig. 2B; see Methods for more details)^24^. Equations in the model were parameterized using maximum growth rates as a function of substrate, half velocity constants, and carrying capacities extracted from the growth data using Logistic or Monod fits (Figs. 2A and S1D and F). Linear models were developed to capture relationships between NO_3_^−^, NO_2_^−^, and N_2_O concentration and maximum OD600 (Fig. S1E) and used in simulations to predict monoculture and SynCom growth kinetics in new experiments. The simulation was optimized to achieve better accuracy by using initial biomass as a free model parameter to effectively adjust lag phase. An initial model formulation for the reduction of NO_3_^−^ to N_2_, combining the reduction of NO_2_^−^ and NO into a single reaction (i.e., reduction of NO_2_^−^ to N_2_O), yielded reasonably accurate prediction of metabolite turnover and growth kinetics for R12 (Fig. S2B). However, the period of maximum growth was overestimated for 3H11 and the carrying capacity (max OD600) achieved by R12 was not well represented. Furthermore, prediction of SynCom dynamics was quite poor (Fig. S2B). While the introduction of NO_2_^−^ inhibition improved predicted growth dynamics of monocultures and SynCom, overall model accuracy was still poor (Fig. 2C, S2C and D). The period of exponential growth was overestimated and NO_2_^−^ turnover was not well predicted.

These results challenged our model assumptions and forced us to reconsider the scheme of metabolite exchange. The most plausible variation was to include exchange of NO, assuming that a significant fraction of the NO produced by R12 is available for reduction by 3H11. This scenario would be possible if the affinity and rate of NO reduction was considerably higher for 3H11 than R12, as was the case for NO_3_^−^. The contribution of the reduction of NO_2_^−^ to N_2_O was subsequently doubled for 3H11 (assuming a similar contribution to its growth) and halved for R12, which was estimated from the energetics of these processes due to absence of measured rates of NO reduction. This assumption significantly improved predicted growth dynamics of the SynCom and offered explanation for the dominance of 3H11 in the community, in contrast to its relatively poor growth in monoculture (Fig. 2C, S2C). Thus, kinetic modeling was essential to revealing the metabolite exchange governing SynCom growth dynamics.

### Community growth drives global changes in transcriptional states of 3H11 and R12

In order to characterize the impact of paired growth on organismal physiology, we compared temporal changes in transcriptomes of each isolate grown as a monoculture and as a SynCom member. Transcriptomes were longitudinally profiled at four time points that corresponded with early-, mid-, and late-phases of growth for all cultures with the exception of the 3H11 monoculture, which was profiled at two time points because of its fast growth rate and limited biomass (Fig. 3A and B). Using DESeq2^25^ to identify differentially expressed genes (DEGs), we discovered that 321 of 4,670 genes in 3H11 and 480 of 3,493 genes in R12 were differentially regulated across different growth phases in the mono-and co-culture contexts (Fig. 3C, Supplementary Table 1). K-means clustering and pathway ontology enrichment analysis (Fig. 3D) identified significant expression changes in a few metabolic pathways, including denitrification, amino acid metabolism, and central carbon metabolism (Fig. 3D and E).

**Figure 3.**
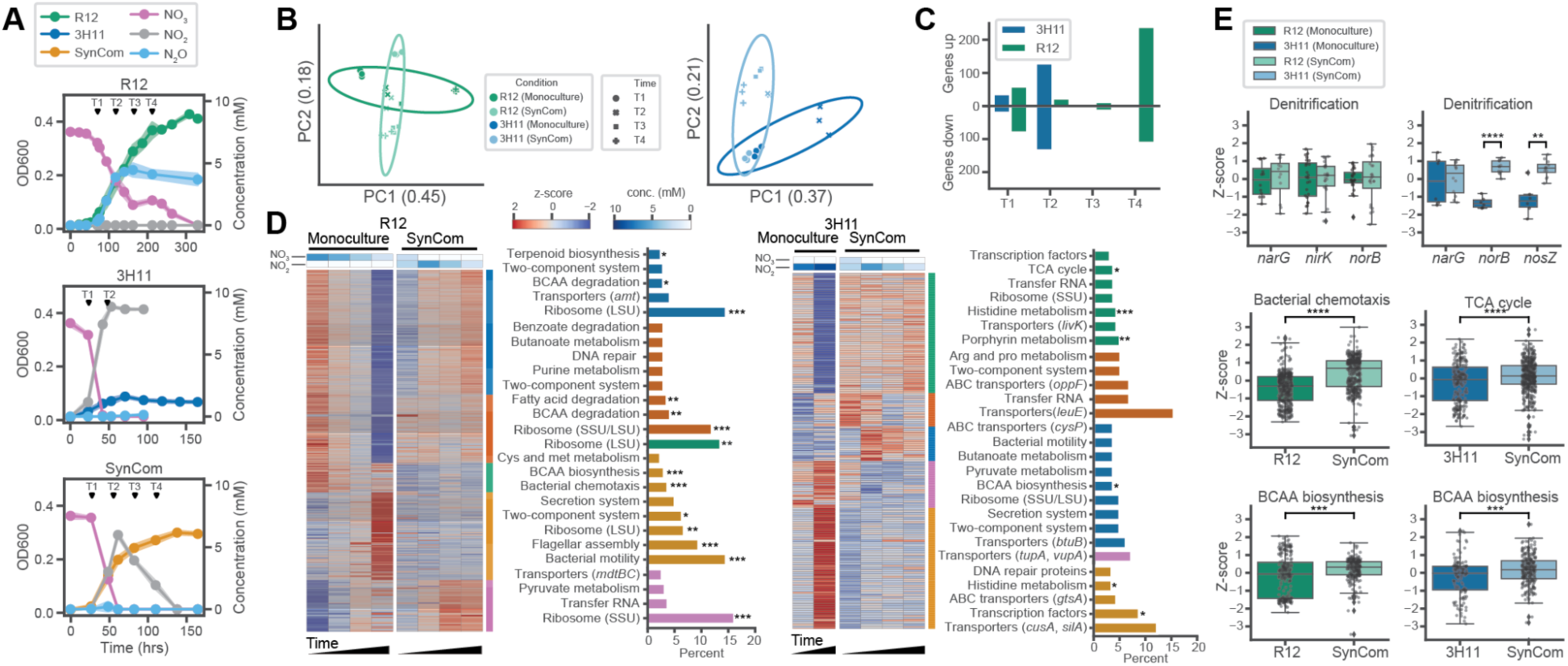
Community context drives changes in the global transcriptional states of 3H11 and R12. (**A**) Growth characteristics of R12, 3H11, the SynCom, and associated transcriptomics sampling points. Black triangles mark condition specific time points (T1-T4). Shading around trend lines represents standard deviations and points represent averages. (**B**) PCA of R12 and 3H11 gene expression across monoculture and co-culture growth contexts. DESeq2 log normalized gene expression Z-scores used. Z-scores computed across growth contexts for R12 and 3H11. R12 and 3H11 specific DEG counts for comparisons across time and growth contexts. (**D**) K-means clustered R12 and 3H11 specific DEGs using k=5. Expression Z-scores are displayed across time and growth contexts. Associated NO_3_^−^ and NO_2_^−^ concentrations are displayed above each heatmap. To the right of each heat map, the most abundant KEGG subpathway terms are displayed for each cluster. Bars are colored by the cluster they represent. For ribosome terms, the most abundant subunit is noted. Large subunit (LSU) or small subunit (SSU). For ABC transporter and transporter terms, a representative gene in each cluster is noted (See Supplementary Table 1 for full list of DEGs and associated clusters). Subpathway terms are noted which were significantly enriched in each cluster using the hypergeometric test. *, *p*<0.05; **, *p*<0.01; ***, *p*<0.001 (**E**). Boxplots of gene and pathway expression Z-scores. Bars indicate comparisons for which differences were significant using Welch’s t-test. *, *p*<0.05; **, *p*<0.01; ***, *p*<0.001

Expression patterns supported the hypothesis that 3H11 is the primary reducer of NO and N_2_O when paired with R12 (Fig 3E). Both 3H11 *norB* and *nosZ* were significantly upregulated in the SynCom context (Welch’s t-test *t*=-11.8, *p*<10^−6^ for *norB*; *t*=-4.5; *p*=0.003 for *nosZ*). In addition to its overall higher average expression in the SynCom context, *norB* was further upregulated during NO_2_^−^ reduction, reaching maximum levels in late-log phase (T3). Transcript levels of 3H11 *nosZ* increased from early-(T1) to mid-log phase (T2) in the SynCom context and decreased thereafter, mirroring corresponding changes in NO_2_^−^ levels (Fig. S3A). In contrast, transcript levels of all R12 denitrification genes were relatively stable across conditions (Fig. 3E), except for *nirK*, which was differentially expressed in the SynCom during the late-log phase of growth (Fig. S3B). Transcript level of *nirK* increased in the SynCom relative to monoculture growth during the period of active NO_2_^−^ reduction (log_2_ fold change=2.62, p<10^−6^ for comparison of T4 *nirK* expression; Fig. S3B). Similarly, R12 *narG* was achieved maximum expression levels earlier (95 hrs) in the SynCom relative to the monoculture context (119 hrs, Fig. 3E, Fig. S3B), which coincided with the timing of NO_2_^−^ accumulation in each of these contexts. This observation was consistent with a higher maximum growth rate and shorter lag phase of R12 monocultures during growth on NO_2_^−^ relative to NO_3_^−^ (Fig. 2A, S1A, S1C) and the increased rate of NO_3_^−^ reduction by the SynCom relative to monocultures (Fig. S2A).

Transcriptional changes across multiple pathways suggested additional mechanisms of competition and cooperation between R12 and 3H11 as a SynCom. For example, expression patterns for 6 of 34 tricarboxylic acid (TCA) cycle genes in 3H11 and 4 of 33 genes in R12 coincided with the timing of NO_3_^−^ and NO/N_2_O reduction across growth contexts (Fig. 3E). Expression of TCA cycle genes in 3H11 coincided with reduction of NO_3_^−^ (T1 in both monoculture and SynCom contexts) and with NO_2_^−^ (T4 in SynCom context) (Fig. S3A). Unlike 3H11, expression of TCA cycle genes in R12 were lower in the initial time points, when NO_3_^−^ was reduced, and increased in later phases of growth, when NO_2_^−^ and N_2_O were being reduced (e.g., R12_3138, *fumA*, downregulated in T1, log_2_ fold change=-2.14, p<10^−6^ and upregulated in T4, log_2_ fold change=3.06, p<10^−6^; Fig S3B). Additionally, 38 motility-associated genes (e.g., *cheA-Z*, *flgA-M*, *fliD-Q*, and *pilA*) were upregulated in R12 in the SynCom context, which may be a mechanism for interaction with 3H11 or competition for NO_3_^−^ (Fig. 3E). Interspecies exchange of amino acids in the SynCom was suggested by the differential regulation of amino acid metabolism genes (74 total in 3H11 and R12). While both species upregulated biosynthesis of branched-chain amino acids (BCAAs) in the SynCom, there was coordinated expression of genes for leucine efflux in 3H11 and upregulation of BCAA degradation genes in R12 (Supplementary Table 1). The coordinated exchange of individual amino acids suggested by the expression data was also consistent with a SynCom metabolic model predicting a 2-3% increase in yield with exchange of individual BCAAs (Fig. S3C). Collectively, the gene expression analysis provided support for the hypothesis that 3H11 is the primary reducer of NO and N_2_O and identified potentially additional mechanisms of metabolic interplay contributing to the improved growth characteristics of the SynCom.

### Proteomics analysis of the SynCom supports the dominant role of 3H11 across varied NO_3_^−^ concentrations and reveals global physiological adaptations

We investigated the consequence of environmental context on the composition and functional dynamics of the SynCom, by performing mass spectrometry-based proteomic profiling across varying initial co-culture conditions, including NO_3_^−^ concentrations (1-40 mM NO_3_^−^, 50% 3H11) and community compositions (10 mM NO_3_^−^, 5-98% 3H11) (Fig. S4). Remarkably, regardless of the ratio of the two organisms in the inoculum, abundance of unique peptides (Fig. S4C) revealed that at steady state the SynCom composition converged to ∼65% 3H11 and ∼35% R12 (Fig. 4A), which was consistent with the kinetic model-predicted dominant role of 3H11 (Fig. 2C)^26^.

**Figure 4.**
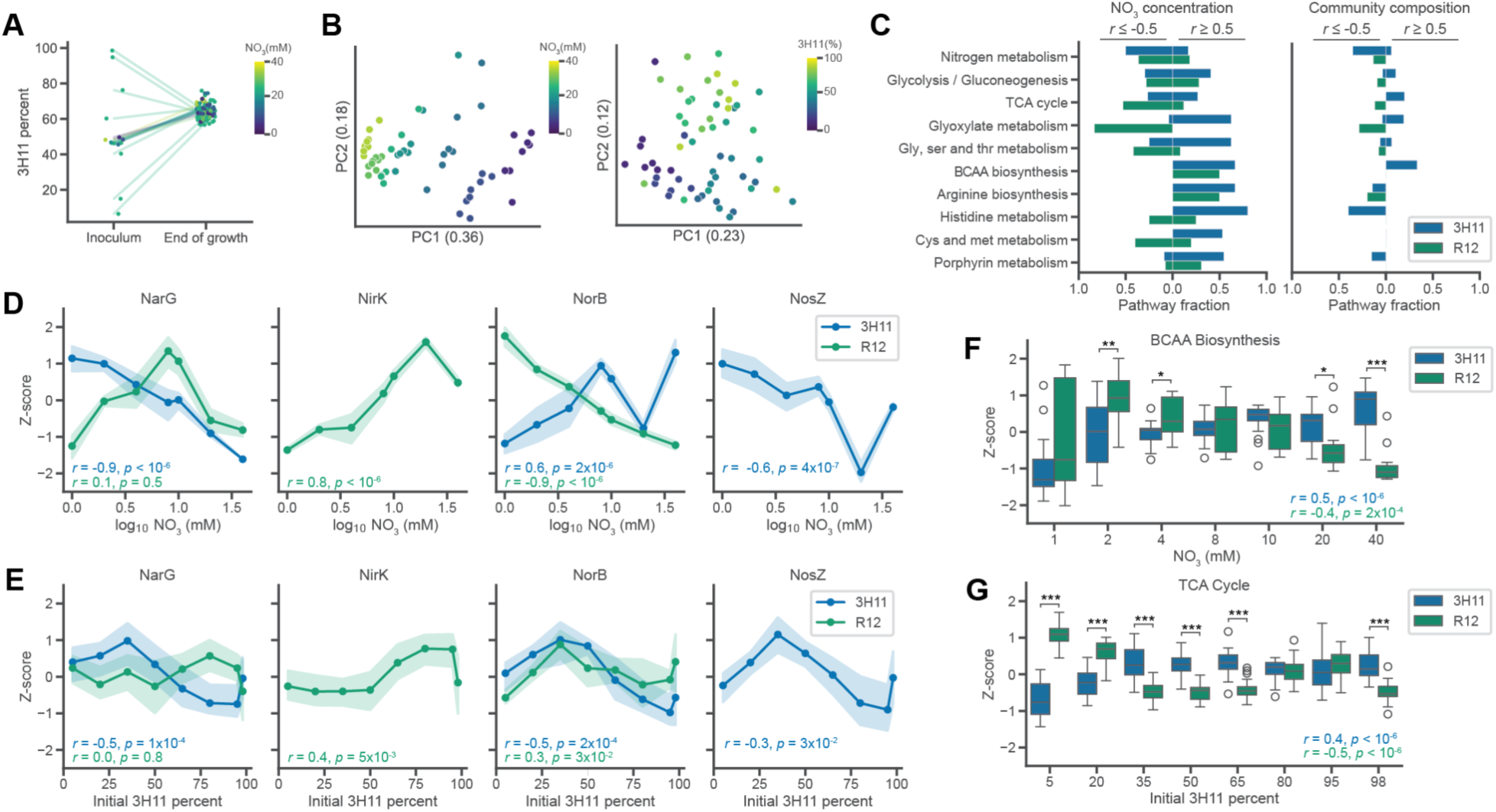
Variation in NO_3_ and community composition support the dominant role of 3H11 in the SynCom. (**A**) 3H11 SynCom percentage across variation in initial community composition and NO_3_ concentration. Community composition measured before inoculation and at the end of growth. Color indicates NO_3_^−^ concentration used and lines connect initial and end points of individual cultures. (**B**) PCA of SynCom end-point proteome Z-scores across variation in initial community composition and NO_3_^−^ concentration. Color used to represent growth conditions. (**C**) Fraction of detected 3H11 and R12 specific pathway proteins significantly correlated with NO_3_^−^ and community composition. (**D**) End-point 3H11 and R12 denitrification protein Z-scores as a function of initial NO_3_^−^ concentration. (**E**) End-point 3H11 and R12 denitrification protein Z-scores as a function of initial community composition. Points represent an average of eight replicates, error bands display standard deviation. 3H11 and R12 specific Pearson *r* and *p* values are displayed on each subplot. (**F**) Boxplots of BCAA biosynthesis protein Z-scores across NO_3_ concentration. (**G**) Boxplots of TCA cycle protein Z-scores across community composition. Bars indicate comparisons for which differences were significant using Welch’s t-test. *, *p*<0.05; **, *p*<0.01; ***, *p*<0.001.

Altogether, 36% of the variance in abundance profiles of 2,401 SynCom proteins was captured by variation in NO_3_^−^ concentration of the growth medium (*r*=-0.94, *p*<10^−6^; Fig. 4B). In fact, abundance changes in nearly half of the proteome (1,105 proteins) were significantly correlated with NO_3_^−^ concentration (*r*≤0.5 or *r*≥0.5 and *q*<0.05; Fig. 4C). Globally, protein abundance changes were also significantly influenced by initial community composition, which showed moderate correlations with PC1 and PC2 (*r*=0.58, *p*<10^−6^; *r*=0.32, *p*=0.011 for PC1 and PC2 respectively; Fig. 4B). Changes in abundance of 206 proteins were significantly correlated with community composition (*r*≤0.5 or *r*≥0.5 and *q*<0.05). Broadly, most proteins correlated with NO_3_^−^ concentration and community compositions were from specific pathways and processes, including energy metabolism, central carbon metabolism, and amino acid metabolism (Fig. 4C).

Abundance patterns in denitrification pathway enzymes suggested that both R12 and 3H11 have distinct regulatory schemes that benefit overall growth characteristics of the SynCom across a range of NO_3_^−^ concentrations (Fig. 4D). For instance, the inverse correlation of 3H11 NarG protein abundance and NO_3_^−^ levels suggests that regulation of this enzyme is highly sensitive to repression by NO_2_^−^. By contrast, expression of R12 NarG was upregulated by low concentrations of NO_3_^−^ (<10mM), but was repressed at concentrations >10mM, also likely due to NO_2_^−^ accumulation. The abundance of R12 NirK was significantly correlated with NO_3_^−^ concentration, suggesting that its expression was activated by and proportional to NO_2_^−^ levels. Together these patterns suggest regulatory mechanisms that balance activities of NarG and NirK to limit the accumulation of NO_2_^−^ in the SynCom. Interestingly, trends in NorB were opposite for R12 and 3H11. Abundance of NorB increased for 3H11 and decreased for R12 as a function of NO_3_^−^. Additionally, 3H11 NosZ abundance decreased with NO_3_^−^ (Fig. 4D). Less pronounced but similar trends were observed for NarG and NirK across variation in community composition (Fig. 4E). The decrease in NorB and NosZ with increase in the initial abundance of 3H11 (Fig. 4E) is likely due to NO_2_^−^ toxicity, since the rate of production of NO_2_^−^ by 3H11 exceeds its reduction by R12. Thus, correspondence in trends between NarG and NirK across conditions suggested that NO_2_^−^ accumulation resulted from an increase in NO_3_^−^ concentration as well as an increase in the proportion of 3H11 in the initial composition of the SynCom.

Protein abundance patterns across multiple pathways showed opposite trends between 3H11 and R12 (Supplementary Table 2). For instance, whereas 40 of the 67 central carbon metabolism enzymes in R12 that were assayed were negatively correlated with NO_3_^−^, 68 of 87 enzymes in the corresponding pathways in 3H11 were positively correlated with NO_3_^−^ (Fig. 4F, Supplementary Table 2). Similar anti-correlated trends in protein abundance changes in 3H11 and R12 were also observed with respect to initial community composition (e.g., proteins of TCA and glyoxylate cycles). Collectively these trends in protein abundance changes suggest that 3H11 outcompeted R12 for substrates (i.e., acetate, NO_3_^−^, NO etc.) with competition for substrates becoming more pronounced with increase in relative proportion of 3H11 in the inoculum, as well as with higher concentrations of NO_3_^−^ (Fig. 4G). Alternatively, these trends may suggest that key metabolic processes, such as amino acid biosynthesis pathways (Supplementary Table 2), were partitioned across the two organisms, complementing each other’s nutritional needs and capabilities. For instance, while four enzymes of methionine biosynthesis in R12 were downregulated with increasing NO_3_^−^ concentration, a methionine importer, MetQ, was upregulated (*r*=0.7, *p*<10^−6^). This regulation in R12 was likely mediated at the transcriptional level by MetJ, a putative repressor of methionine biosynthesis, which was upregulated with increased NO_3_^−^ (*r*=0.53, *p*=5.6×10^−5^). By contrast, enzymes of methionine biosynthesis in 3H11 were upregulated with increased NO_3_^−^ concentration (*r*>0.5, *p*<10^−6^ for MetE, MetH and MetY). These trends suggested that methionine biosynthesis in 3H11 was complementing the nutritional need of this amino acid in R12. The NO_3_^−^ -induced shifts in abundance of enzymes of amino acid metabolism in both R12 and 3H11 was consistent with corresponding mRNA level changes indicating that much of this regulation was mediated at the transcriptional level. Thus, the proteomics analysis supported conclusions from transcriptomics analysis regarding the role of process partitioning and interplay of varied metabolic processes, including denitrification, central carbon metabolism, and amino acid metabolism, in enhancing growth characteristics of the SynCom.

### Nitrite accumulation drives nitrous oxide emissions across variation in initial nitrate concentration and community composition

We investigated how variation in abiotic factors (specifically, NO_3_^−^ concentration) and biotic properties (specifically, community composition) impact N_2_O emissions. We measured the steady state levels of N_2_O in SynCom cultures across a range of initial NO_3_^−^ concentrations (1 mM vs 40 mM) and community composition (3H11:R12 ratio of 5%:95% vs 98%:2%). We expected that extremes in community composition and NO_3_^−^ concentration might result in differences in community phenotype and N_2_O production/consumption. We anticipated N_2_O accumulation at higher concentrations of NO_3_^−^. At high NO_3_^−^ concentrations acetate would be exhausted before NO_3_^−^ could be completely reduced to N_2,_ given that earlier steps in the pathway (e.g., NO_3_^−^ and NO_2_^−^ reduction) are prioritized by the SynCom. Consistent with this hypothesis, N_2_O accumulation was proportional to initial NO ^−^ However, the observed accumulation of N_2_O at 20 mM NO_3_ ^−^ was well below the 30 mM threshold predicted by stoichiometry to become limiting. In addition, we were surprised to find that N_2_O also increased with increasing proportions of 3H11 in the inoculum, despite its role as the primary N_2_O reducer in the SynCom (Fig. 5A). Together these findings suggested a more complex interplay between 3H11 and R12 underlies the dynamics of denitrification by the SynCom.

**Figure 5.**
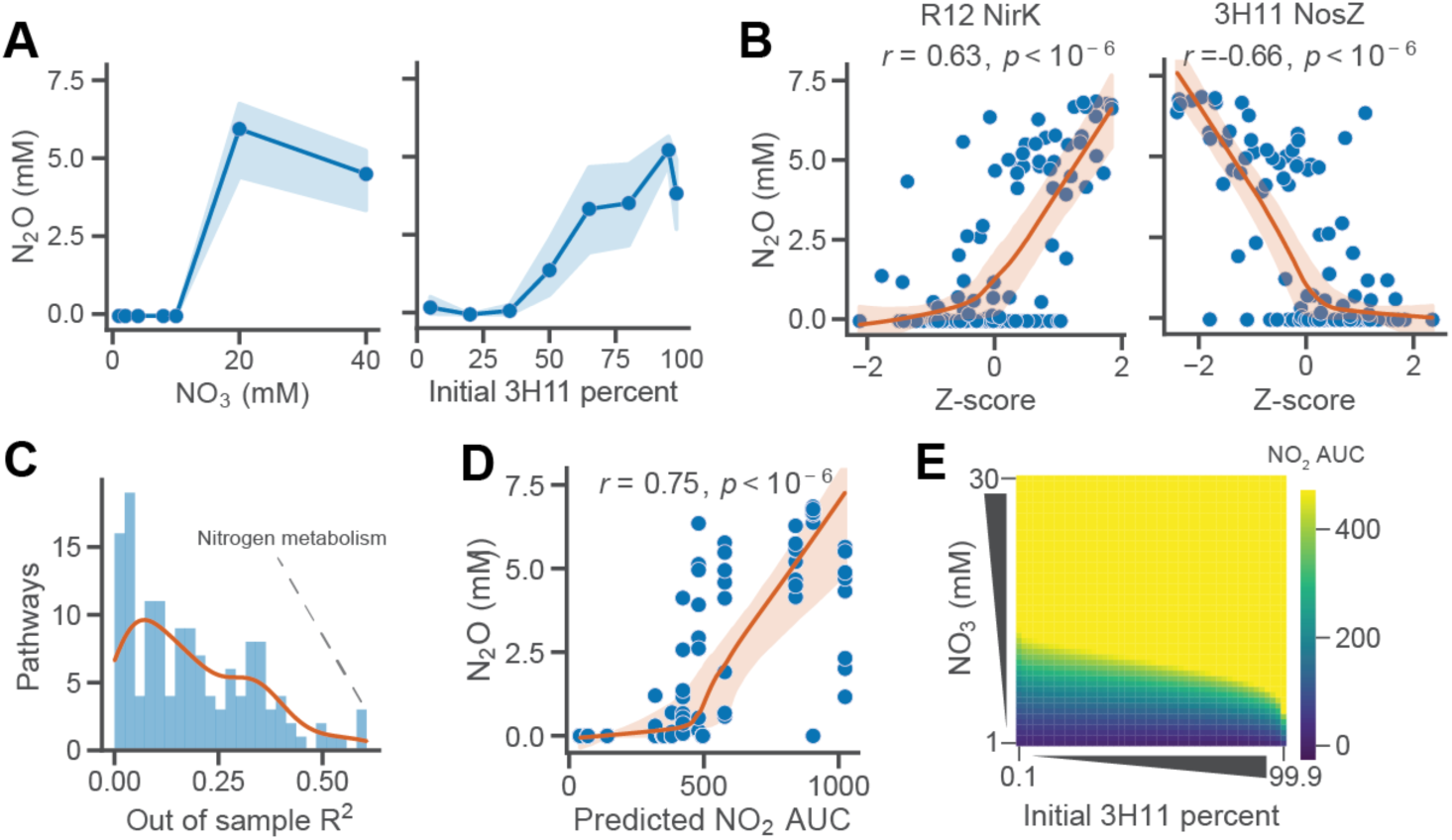
NO_2_^−^ inhibition drives N_2_O accumulation in the SynCom. (**A**) End-point SynCom N_2_O concentration as a function of initial NO_3_^−^ concentration and community composition. Points represent an average of eight replicates, error bands display standard deviation. (**B**) End-point SynCom N_2_O concentration as a function of denitrification protein Z-scores across conditions. Lowess trendline and 95% confidence interval displayed. Pearson *r* and *p* values displayed. (**C**) Distribution of predicted N_2_O out of sample R^2^ using pathway level multi-linear regression. R^2^ of N metabolism proteins noted. (**D**) End-point SynCom N_2_O concentration as a function of predicted SynCom NO_2_^−^ AUC concentration across conditions. Lowess trendline and OLS linear fit displayed with respective 95% confidence intervals. Pearson *r* and *p* values displayed. (**E**) Heatmap of predicted SynCom NO_2_^−^ AUC across variation in initial NO_3_^−^ concentration and community composition.

To characterize possible mechanisms responsible for accumulation of N_2_O by the SynCom, we examined associations between N_2_O concentration and protein abundance across all conditions. We identified 13 proteins that were positively correlated with N_2_O levels across NO_3_^−^ concentrations and community composition (*r*≥0.5 and *q*<0.05) and 12 proteins with abundance changes that were negatively correlated (*r*≤-0.5 and *q*<0.05). Proteins that were positively correlated with N_2_O were largely from R12 (10 of 13) and of unknown function (6 of 10). The four R12 proteins with predicted functions were NirK, an iron transporter, Bfr (bacterioferritin), and Ahr (alcohol/geraniol dehydrogenase). The three 3H11 proteins that were positively correlated with N_2_O were associated with central carbon and amino acid metabolism and included AceB, GlcB (malate synthase), Ddl (D-alanine-D-alanine ligase), and ilvH, ilvN (acetolactate synthase). In contrast, proteins with abundance changes negatively correlated with N_2_O levels were primarily from 3H11 (10 of 12) and related to energy metabolism (7 of 10). Two of the seven energy metabolism proteins were cytochromes, while the rest were from the denitrification pathway, including two copies of NarG, NarH, NosZ, NosR (a transcriptional regulator of NosZ) and PhasZ (PHB depolymerase; Fig. 5B).

Using a multi-linear regression framework, we investigated the ability of protein abundances within specific pathways to predict observed final N_2_O levels (Fig. 5C, see Methods for additional details). As expected, protein abundances within the N metabolism pathway had the best performance in predicting final N_2_O levels (out of sample R^2^=0.53). The patterns of correlations between the abundance of denitrification enzymes in the two organisms and N_2_O (e.g., positive correlation of R12 NirK and negative correlation of 3H11 NosZ), suggested that NO_2_^−^ toxicity might be the driver of N_2_O emissions in the SynCom. This hypothesis was supported by the strong correlation between both maximum concentration and cumulative (i.e., area under the curve (AUC)) levels of NO_2_^−^ predicted by the kinetic model, and the observed final N_2_O levels for each experimental context (i.e., initial NO_3_^−^ levels and community composition). Relationships between final N_2_O levels and both predicted NO_2_^−^ metrics identified an inflection point which likely corresponds to the concentration at which NO_2_^−^ became toxic, inhibiting process partitioning between the two organisms, which ultimately manifested in increased N_2_O emissions (Fig. 5D). This inflection point for N_2_O production occurred at ∼7.5 mM NO_2_^−^, which was in line with the observation that growth of 3H11 monocultures were completely inhibited at NO_2_^−^ concentrations >5 mM (Fig. 2A, Fig. S1).

Finally, we investigated factors governing NO_2_^−^ accumulation by using the SynCom kinetic model. We predicted the total amount of NO_2_^−^ produced by the SynCom across a range of initial co-culture conditions, community compositions (0.1-99.9 % 3H11) and NO_3_^−^ concentrations (1-30 mM). This analysis revealed that NO_3_^−^ concentration and community composition had non-linear effects on the accumulation of NO_2_^−^ to a level that would result in N_2_O off gassing through growth inhibition of 3H11. Together our findings demonstrate that during process partitioning of denitrification, N_2_O production (“off gassing”) is primarily driven by NO_2_^−^ toxicity, which manifests from a complex interplay of environmental context and community composition.

## DISCUSSION

Our findings demonstrate how two ecologically relevant partial denitrifiers cooperate as a SynCom to perform complete denitrification (Fig.1) and how differences in mechanisms of pathway partitioning (i.e., exchange of specific metabolites) and kinetics of substrate utilization by individual members can manifest in distinct emergent phenotypes of a microbial community (Fig. 2). Specifically, we discovered four key mechanisms that contributed to the overall improvement in growth dynamics of the SynCom. First, the fast rate of 3H11-mediated reduction of NO_3_^−^ into NO_2_^−^ promoted growth of R12, a finding supported by evidence that monocultures of R12 grew better on NO_2_^−^, relative to NO_3_^−^, with a shorter lag phase and increased growth rate (Fig. 2A and S1). Second, by consuming NO_2_^−^, R12 facilitated increased biomass of 3H11, based on evidence that 3H11 was severely growth inhibited due to accumulation of NO_2_^−^ when grown as a monoculture (Fig. S1), but not in a SynCom context wherein NO_2_^−^ levels increased transiently and returned to undetectable levels (Fig. 2C and S4E). Third, the exchange of NO (instead of N_2_O) from R12 to 3H11 turned out to be a critical model-enabled discovery, which was supported by the subsequent observation that 3H11 *norB* was significantly upregulated in the SynCom context (Fig. 3E). Finally, the finding that N_2_O can itself serve as a source of energy and biomass for 3H11 added evidence for a fourth mechanism for synergistic improvement in specific growth rate of the SynCom (Fig. S1C). Collectively, the four mechanisms incorporated into a kinetic model explained how interplay between the two isolates improved overall growth characteristics of the SynCom (Fig. 2C).

While we note the natural variation in denitrification kinetics that can be achieved by environmental isolates and communities, our findings are consistent with the principle of inter-enzyme competition described by Lilja *et al*.^19^. The observed increase in specific growth rate as well as the increased rate of NO_3_^−^ reduction of the SynCom relative to R12 and 3H11 monocultures (Fig. 1E, F, S3C), suggests that inter-enzyme competition was reduced in the SynCom context. Notably, we observed that *Rhodanobacter* spp. R12 monocultures do not accumulate NO_2_^−^ and can grow with significantly reduced lag time and higher growth rates on NO_2_^−^ relative to NO_3_^−^ (Figs. 1E and 2A). This is in contrast to the strain of *Pseudomonas stutzeri* used by Lilja *et al*., which favors NO_3_^−^ reduction and thus accumulates NO_2_^− 19^. This suggests that the R12 *Rhodanobacter* has evolved a balance between NarG and NirK, or between these enzymes and other rate limiting steps such as NO_3_^−^ transport, that favors reduction of NO_2_^−^ over NO_3_^−^.

*Rhodanobacter spp.* dominate microbial communities within groundwater associated with low pH and high NO_3_^−^ in regions of the ORR, including the specific site where R12 was isolated^8,23,27^. Thus, the groundwater conditions at ORR might provide a selection pressure that favors the balance in NO_3_^−^ and NO_2_^−^ reduction achieved by R12 and possibly other *Rhodanobacter spp*. However, we cannot rule out the possibility that the observed improvements in specific growth rate and NO_3_^−^ reduction rate was due to the alleviation of NO_2_^−^ toxicity. Indeed, N_2_O off gassing by the SynCom in certain contexts (i.e., high proportion of 3H11 and high NO_3_^−^) was largely attributed to the accumulation of NO_2_^−^ due to the increased rate of NO_3_^−^ reduction facilitated by 3H11, which effectively shifted the balance achieved by R12 toward NO_3_^−^ reduction (S3C). Thus, in addition to inter-enzyme competition, natural variation in denitrification kinetics of individual organisms and toxicity of intermediates are important factors that significantly influence composition and functional interactions within a microbial community across environmental contexts.

Our recognition of NO_2_^−^ accumulation as an important driver of N_2_O emissions also offers mechanistic understanding to environmental studies linking NO_2_^−^ with N_2_O emissions. Maharjan and Ventera showed that NO_2_^−^ accumulation was a major driver of N_2_O emissions across different N fertilizer regimes in an agricultural setting and found that mitigation of NO_2_^−^ accumulation reduced N_2_O emissions^28^. NO_2_^−^ has also been implicated in N_2_O emissions in the context of wastewater treatment, where free nitrous acid (HNO_2_) has been shown to drive emissions for denitrifying-enhanced biological phosphorus removal sludge via inhibition of N_2_O reduction^29^. In this context the authors point toward free HNO_2_ rather than NO_2_^−^ itself as the source of inhibition, thus implicating pH as an important factor for the link between NO_2_^−^ and N_2_O emissions in denitrifying communities. Therefore, while the inhibitory effects of NO_2_^−^ are well characterized and have been associated with N_2_O emission in other contexts, here using the 3H11-R12 SynCom we have uncovered how factors, such as NO_3_^−^ concentration, community composition, and substrate utilization kinetics, modulate partitioning of denitrification across a microbial community to drive NO_2_^−^ accumulation and N_2_O emissions.

While the kinetic model provided a mechanistic explanation for improved growth characteristics of the SynCom, it also suggested that functional interactions between the two organisms extended beyond the core process of denitrification. Indeed, gene expression and proteomics analyses uncovered widespread complementary physiological changes across both organisms when they were grown as a SynCom (Figs. 3 and 4). Correlated protein level changes of enzymes of BCAA and histidine biosynthesis in 3H11 with corresponding changes in NO_3_^−^ and N_2_O levels suggested that upregulation of these pathways was likely governed by the availability of NO produced by R12. Upregulation of BCAA biosynthesis may contribute towards maintaining intracellular redox balance for 3H11 and R12, as reported previously in the purple non sulfur bacterium *Rhodospirillum rubrum* growing under photoheterotrophic conditions^30,31^. Additionally, upregulation of leucine export by 3H11 and BCAA degradation genes by R12 (e.g., leucine dehydrogenase, and propionyl-CoA carboxylase) suggested exchange of leucine from 3H11 to R12. Similar trends in protein level changes were observed which suggested that methionine biosynthesis by 3H11 was likely leveraged by R12 through upregulation of uptake and degradation pathways. These observations collectively suggest that when growing in a SynCom context, R12 derives some of its nutritional needs from 3H11, which may be facilitated by coordinated upregulation of flagellar genes^32^. Consistent with these findings, simulations based on the stoichiometric constraints-based metabolic model of the SynCom predicted that exchange of intermediates of central C metabolism and amino acids could contribute up to 10% increase in overall biomass of the SynCom (Fig. S3C). In sum, multiomics profiling and model simulations suggested that synergistic improvement in growth characteristics of the SynCom was likely an outcome of process partitioning across both denitrification as well as other metabolic pathways, explaining why metabolite exchange is ubiquitous across natural microbial communities ^33^. Future work should integrate kinetics of denitrification into metabolic models of monocultures and SynComs to better understand the dynamics of community metabolism and how it changes with growth context^34,35^. This may allow us to better understand how changes in pH, temperature, dissolved oxygen, and other environmentally relevant factors influence cellular physiology, community cooperativity, and the production of N_2_O in natural ecosystems.

## METHODS

### ORR and IMG isolate denitrification pathway composition analysis

Annotated ORR isolate genomes were obtained from KBase (https://narrative.kbase.us/narrative/55476). Additional soil isolate genomes were obtained from the Joint Genome Institute’s (JGI’s) Integrated Microbial Genomes & Microbiomes (IMG/M) database were. The presence of the denitrification genes Nar/Nap, Nir, Nor, and Nos were characterized using KEGG (Kyoto Encyclopedia of Genes and Genomes) Orthology (KO) terms. Isolates with one or more denitrification genes were considered across both datasets. Denitrification pathway composition of the isolates was tabulated using gene presence/absence.

### ORR groundwater genus abundance and chemistry analysis

Groundwater OTU abundance, taxonomy, and chemical data previously described by Smith et al. 2015 were obtained and re-analyzed ^20^. OTU relative abundances were summarized at the genus level and log_10_ transformed. Chemical concentrations were converted to mM and log_10_ transformed where appropriate. Genus presence/absence was determined by examining the overall distribution of log_10_ transformed genus relative abundances and setting a threshold. Genera with a log_10_ abundance > -6 (e.g., relative abundance > 10^−6^) were considered present. This threshold separated the peaks of the bi-model log_10_ genus relative abundance distribution. Statistical analyses, including linear regression, and significance testing, were performed using tools from the python Scikit-Learn, SciPy, and NumPy packages ^36–38^.

### Isolate and SynCom and metabolic models

The genome-scale metabolic models (GEMs) used in this study were reconstructed using the ModelSEED v2 ^39^ app in The Department of Energy (DOE)’s Systems Biology Knowledgebase (KBase) ^40^. Flux balance analysis (FBA) ^41^ was used to perform growth simulations for individual models for R12 and 3H11 and species interactions in the SynCom. The biomass objective function of the individual models was adjusted as needed to represent the abundance of each organism in the SynCom. When simulating community interactions, we limited the exchange of compounds between organisms to the compounds present in the *in silico* growth media.

### Strains and medium preparation

*Rhodanobacter sp. FW510-R12* (R12) and *Acidovorax sp. GW101-3H11* (3H11) were both isolated from contaminated wells at the Oak Ridge Integrated Field Research Center (ORR) as reported in Hemme et al. 2016 and Price et al. 2018, respectively ^42,43^. NO_3_ reduction growth studies were performed in Balch tubes (10-mL culture volume) at 30°C in a defined minimal medium as described previously containing 20 mM sodium acetate and 10 mM sodium NO_3_ at pH 7.2 with a 80:20 N_2_-CO_2_ headspace ^9^. A small amount of yeast extract (0.1 g/L) was added to the medium before autoclaving to support growth of 3H11 and R12. For experiments that characterized growth on variable amounts of NO_3_, NO_2_^−^, and N_2_O media was first prepared in serum bottles. Vitamin and phosphate solutions were added to serum bottle media and then aliquoted into N_2_-CO_2_ flushed and pressurized Balch tubes (10 mL each, ∼1 psi overpressure). For experiments characterizing growth of 3H11 on N_2_O, injections of pure N_2_O (0-2.59 mL at 14.5 psi overpressure) into Balch tubes were performed to achieve desired concentrations (0-20 mM).

### Preparation of R12 and 3H11 co-cultures and pure cultures

From freezer stocks, cells were plated on R2A agar and incubated at 30 °C. Following growth of isolated colonies, colonies were picked and used for inoculation of R2A liquid medium cultures. Liquid R2A cultures were incubated at 30°C with shaking at 100 rpm. Following overnight growth, optical density at 600 nm (OD600) was measured and cell density was normalized to using R2A liquid medium if necessary. With starting cultures at ∼0.5 OD600 units, 100 uL of *Rhodanobacter sp. FW510-R12* (R12) and 100 uL of *Acidovorax GW101-3H11* (3H11) were inoculated into a NO_3_ reduction medium to establish co-cultures. For pure cultures 200 uL of either strain was utilized to normalize starting cell densities. Initial cell densities were normalized to ∼0.01 OD600 units across all experiments and conditions. Biological replicates varied from 3-8 across experiments. For all experiments cell concentration was monitored in Balch tubes with periodic measurements of the optical density. Initial experiments (Figs. 1E, 2C, and 3A) measured optical density 1-3 times daily using a Spectronic 200 spectrophotometer (Thermo Fisher). Later experiments (Figs. 2A, 4, and 5) leverage an automated OD600 measurement with shaking at 100 rpm previously described ^44^. In this context, measurements were taken every 5-30 minutes and instrument voltage was converted to OD600 by calibration. Sample OD600 was measured before and after each experiment to calibrate individual sample voltage ranges.

### Measurement of N species

NO_3_, NO_2_^−^, and acetate concentrations were analyzed using the ion-chromatography Dionex ICS-5000 system with the IonPac® ICE-AS6 column (Thermo Fisher). Medium N_2_O concentration was quantified by sampling 200 uL medium into GC vials containing 200 uL 8% PFA in 1x PBS (to achieve a final concentration of 4% PFA). Vials were allowed to equilibrate and headspace N_2_O was measured using a Shimadzu GC-2014 with an AOC-6000 autosampler. Vials were equilibrated for 60 seconds in a 35 ℃ oven with shaking, and then a 1000 μL sample was aspirated from the headspace of the vial with a syringe preheated to 100 ℃. Run conditions were as follows: inlet temperature 325 ℃, oven temperature 80 ℃, column flow rate 20 mL/min, TCD temperature: 150 ℃, current: 80 mA, ECD temperature: 325 ℃, current: 2 nA, run time: 20 minutes, carrier gas: argon, makeup gas: N. GC vial headspace N_2_O ppm values were converted to culture vessel liquid N_2_O concentration (mM) using Henry’s law. Total culture vessel N_2_O concentration reported was obtained using mass balance and assuming equilibrium.

### Monoculture and SynCom kinetic models

Kinetic models were developed using a modified Monod framework by integrating Logistic representation of carrying capacity into equations describing growth kinetics as a function of metabolite concentrations ^24^. Equations in the model were of the form 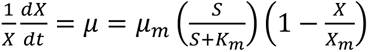, where *X* is cell population size, *S* is substrate concentration, *K_m_* and *μ_m_* are substrate specific Monod kinetic parameters, and *X_m_* is cell specific capacity. Substrate specific rates were assumed to be additive (e.g., *μ_R_*_12_ = *μ_N_*_03_ + *μ_N_*_02_ + *μ_N_*_0_). Equations were parameterized using maximum growth rates as a function of substrate, half velocity constants, and carrying capacities extracted from species specific growth data using Logistic and Monod fits. To represent carrying capacity as a function of substrate concentration, linear models were developed to capture relationships between NO_3_^−^, NO_2_^−^, N_2_O concentration and maximum OD600. Carrying capacity models were of the form *OD_R_*_12_ = *β_N_*_03_*X_N_*_03_ + *β_N_*_02_*X_N_*_02_ and *OD*_3*H*11_ = *β_N_*_03_*X_N_*_03_ + *β_N_*_02_*X_N_*_02_ + *β_N_*_20_*X_N_*_20_ for R12 and 3H11 respectively. Linear model parameters were estimated by OLS fit of monoculture maximum OD600 and initial substrate concentration data. Kinetic model simulations were performed using the python package Tellurium ^45^. Statistical analyses, including linear regression, and significance testing, were performed using tools from the python Scikit-Learn, SciPy, and NumPy packages ^36–38^.

### Transcriptomics profiling and analysis

#### (i) Sample collection and sequencing

R12-3H11 co-cultures and pure cultures were harvested for transcriptome profiling in biological triplicates across time points (Fig. 3). Cell pellets were harvested by centrifugation at 4,000 × g and flash frozen in liquid N_2_. Total RNA was extracted using the MasterPure™ Complete DNA and RNA Purification Kit from Epicentre. All samples were treated with Invitrogen TURBO DNA-Free kit to remove DNA contamination. The Illumina Stranded Total RNA Prep Ligation with Ribo-Zero Plus (Illumina) was used for rRNA depletion and library preparation. Sequencing was performed using the NextSeq 500 platform (2 by 75 bp, Illumina) with 10 to 15 million reads per sample.

#### (ii) Read processing

RNA sequencing reads were analyzed with FastQC according to Illumina’s default quality filtering process and then trimmed using base quality scores by Trimmomatic ^46,47^. A quality score of 20 was used for read trimming and quality filtering. The abundance of 3H11 and R12 specific transcripts were quantified using the concatenated genomes of *Rhodanobacter sp. FW510-R12* (NCBI BioProject PRJNA255897) and *Acidovorax sp. GW101-3H11* (NCBI BioProject PRJNA314893) and Kallisto ^48^.

#### (iii) Differential expression analysis, clustering, and functional enrichment

Normalized expression data and differentially expressed genes (DEGs) were obtained from DESeq2 ^25^ using pairwise comparisons of growth phase matched monoculture and SynCom transcriptomes. Genes were reported as dysregulated if the log2 fold change magnitude ≥ 2 and significant (*p* < 0.05 and Benjamini-Hochberg false discovery rate *q*<0.01). DEGs were clustered using the scikit-learn implementation of the k-means algorithm ^36^. Statistical analyses, including principal-component analysis (PCA), significance testing, and functional enrichment were performed using tools from the Python Scikit-learn, SciPy, and NumPy packages ^36–38^. The significance of KEGG subpathway term enrichment among k-means clusters was assessed by comparing term frequencies within each cluster to their frequencies in the genome using the hypergeometric test ^49^. Significantly enriched terms (*p*<0.05 and Benjamini-Hochberg false-discovery rate *q*<0.01) were reported ^50^. KEGG pathway level expression comparisons were assessed using DESeq2 normalized gene expression Z-scores and Welch’s t-test.

### Proteomics profiling and analysis

#### (i) Sample collection and peptide quantification

R12-3H11 co-cultures and pure cultures were harvested prior to inoculation and at the end of growth (stationary phase) for proteomics profiling using 8 biological replicates (Fig. 4, S4). Cell pellets were harvested by centrifugation at 4,000 × g and flash frozen in liquid N_2_ and stored at -80 °C until further processing. Protein was extracted from cell pellets and tryptic peptides were prepared by following an established proteomic sample preparation protocol ^51^. Briefly, cell pellets were resuspended in Qiagen P2 Lysis Buffer (Qiagen, Germany) to promote cell lysis. Proteins were precipitated with addition of 1 mM NaCl and 4 x vol acetone, followed by two additional washes with 80% acetone in water. The recovered protein pellet was homogenized by mixing with 100 mM ammonium bicarbonate in 20% methanol. Protein concentration was determined by the DC protein assay (BioRad, USA). Protein reduction was accomplished using 5 mM tris 2-(carboxyethyl)phosphine (TCEP) for 30 min at room temperature, and alkylation was performed with 10 mM iodoacetamide (IAM; final concentration) for 30 min at room temperature in the dark. Overnight digestion with trypsin was accomplished with a 1:50 trypsin:total protein ratio. The resulting peptide samples were analyzed on an Agilent 1290 UHPLC system coupled to a Thermo Scientific Orbitrap Exploris 480 mass spectrometer for discovery proteomics ^52^. Briefly, peptide samples were loaded onto an Ascentis® ES-C18 Column (Sigma–Aldrich, USA) and were eluted from the column by using a 10 minute gradient from 98% solvent A (0.1 % FA in water) and 2% solvent B (0.1% FA in ACN) to 65% solvent A and 35% solvent B. Eluting peptides were introduced to the mass spectrometer operating in positive-ion mode and were measured in data-independent acquisition (DIA) mode with a duty cycle of 3 survey scans from m/z 380 to m/z 985 and 45 MS2 scans with precursor isolation width of 13.5 m/z to cover the mass range. DIA raw data files were analyzed by an integrated software suite DIA-NN ^53^. The database used in the DIA-NN search (library-free mode) leveraged the combined 3H11 and R12 annotated proteome FASTA sequences plus the protein sequences of common proteomic contaminants. DIA-NN determines mass tolerances automatically based on first pass analysis of the samples with automated determination of optimal mass accuracies. The retention time extraction window was determined individually for all MS runs analyzed via the automated optimization procedure implemented in DIA-NN. Protein inference was enabled, and the quantification strategy was set to Robust LC=High Accuracy. Output main DIA-NN reports were filtered with a global FDR=0.01 on both the precursor level and protein group level.

#### (ii) Species relative abundance estimates

The quantification of species biomass contributions in the community was based on the input amounts of the sum of peptide ion intensities using only unique peptides from each species. This method has been benchmarked previously as one of the three proteomic quantification methods for accurate estimation of species-level biomass contribution in microbial communities ^26^.

#### (iii) Statistical analyses

Statistical analyses, including PCA, linear regression, and significance testing, were performed using tools from the Python Scikit-learn, SciPy, and NumPy packages ^36–38^. The significance of individual protein abundance changes was assessed by correlating protein abundance with initial NO_3_^−^ levels, inoculum 3H11 percentage, and steady state N_2_O levels. Pearson *r* values were used to identify significant hits, proteins with *r*≤0.5 or *r*≥0.5 and *q*<0.05(Benjamini-Hochberg false-discovery rate correction) were considered ^50^. KEGG subpathway level protein abundance correlations with steady state N_2_O levels were performed using data randomly split into training (70% of samples) and testing (30% samples) sets. KEGG subpathway terms were used to identify groups of proteins and multi-linear regression was performed with the training set. Model accuracy was then assessed using the test set and out of sample R^2^ was reported.

## DATA AVAILABILITY

Data summary tables (e.g., growth and metabolite concentration data, processed transcriptomics and proteomics data, and metabolic model outputs) and code used for data analysis and figure generation can be found at Github at https://github.com/baliga-lab/3H11_R12_SynCom. Raw sequencing data used for the transcriptomics data was uploaded to NCBI GEO repository with the dataset identifier GSE272493. The generated mass spectrometry proteomics data have been deposited to the ProteomeXchange Consortium via the PRIDE partner repository with the dataset identifier PXD051979 ^54^.

## ACKNOWLEDGMENTS

This work was supported by ENIGMA-Ecosystems and Networks Integrated with Genes and Molecular Assemblies (http://enigma.lbl.gov), a Scientific Focus Area Program at Lawrence Berkeley National Laboratory based upon work supported by the U.S. Department of Energy, Office of Science, Office of Biological & Environmental Research under contract number DE-AC02-05CH11231. This work was also supported as part of the Genomic Sciences Program. The DOE Systems Biology Knowledgebase (KBase) is funded by the U.S. Department of Energy, Office of Science, Office of Biological and Environmental Research under Award Numbers DE-AC02-05CH11231, DE-AC02-06CH11357, DE-AC05-00OR22725, and DE-AC02-98CH10886.

---

**Supplemental figure 1.**
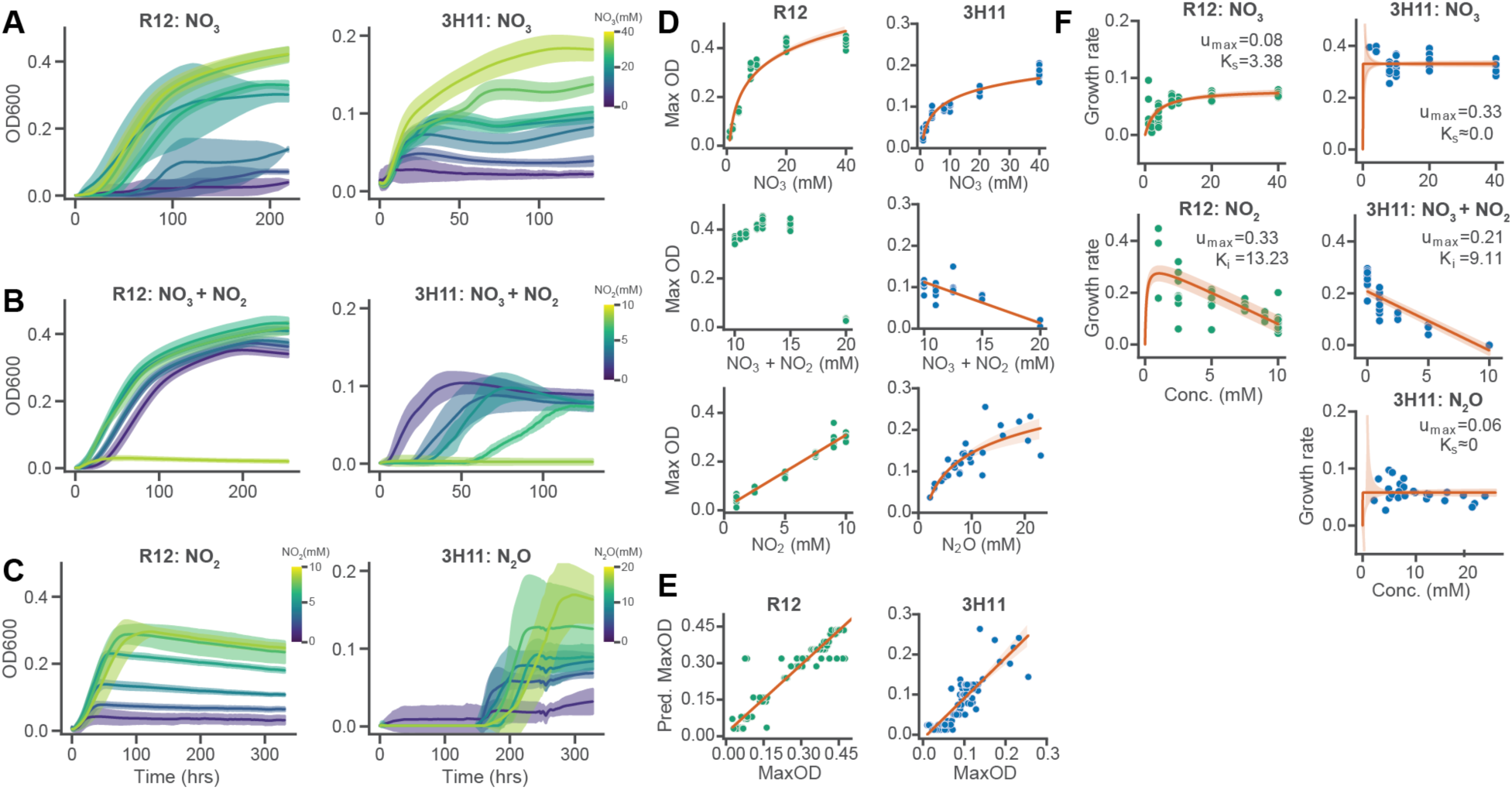
Growth characteristics of 3H11 and R12 across variation in substrate concentration. (**A**) Growth of R12 and 3H11 across variation in initial NO_3_ concentration. Trendlines display average trajectories across eight replicates and error bands display standard deviations. (**B**) Growth of R12 and 3H11 across variation in initial NO_2_ concentration supplemented with 10 mM NO_3_. (**C**) Growth of R12 and 3H11 across variation in initial NO_2-_ and N_2_O concentration respectively. Trendlines display average trajectories across eight replicates and error bands display standard deviations. Trendlines are colored by respective substrate concentrations. (**D**) Maximum OD600 achieved by 3H11 and R12 across growth conditions as a function of substrate concentration. OLS linear or logarithmic fits to data are displayed with 95% confidence intervals where appropriate. (**E**) Predicted maximum OD600 vs measured maximum OD600 using multi-linear fit to linear portions of substrate vs maximum OD600 data. (**F**) Maximum growth rate as a function of substrate concentration across conditions for 3H11 and R12. Fits of Monod substrate kinetics or Monod product inhibition kinetics displayed with 95% confidence intervals and associated parameters.

**Supplemental figure 2.**
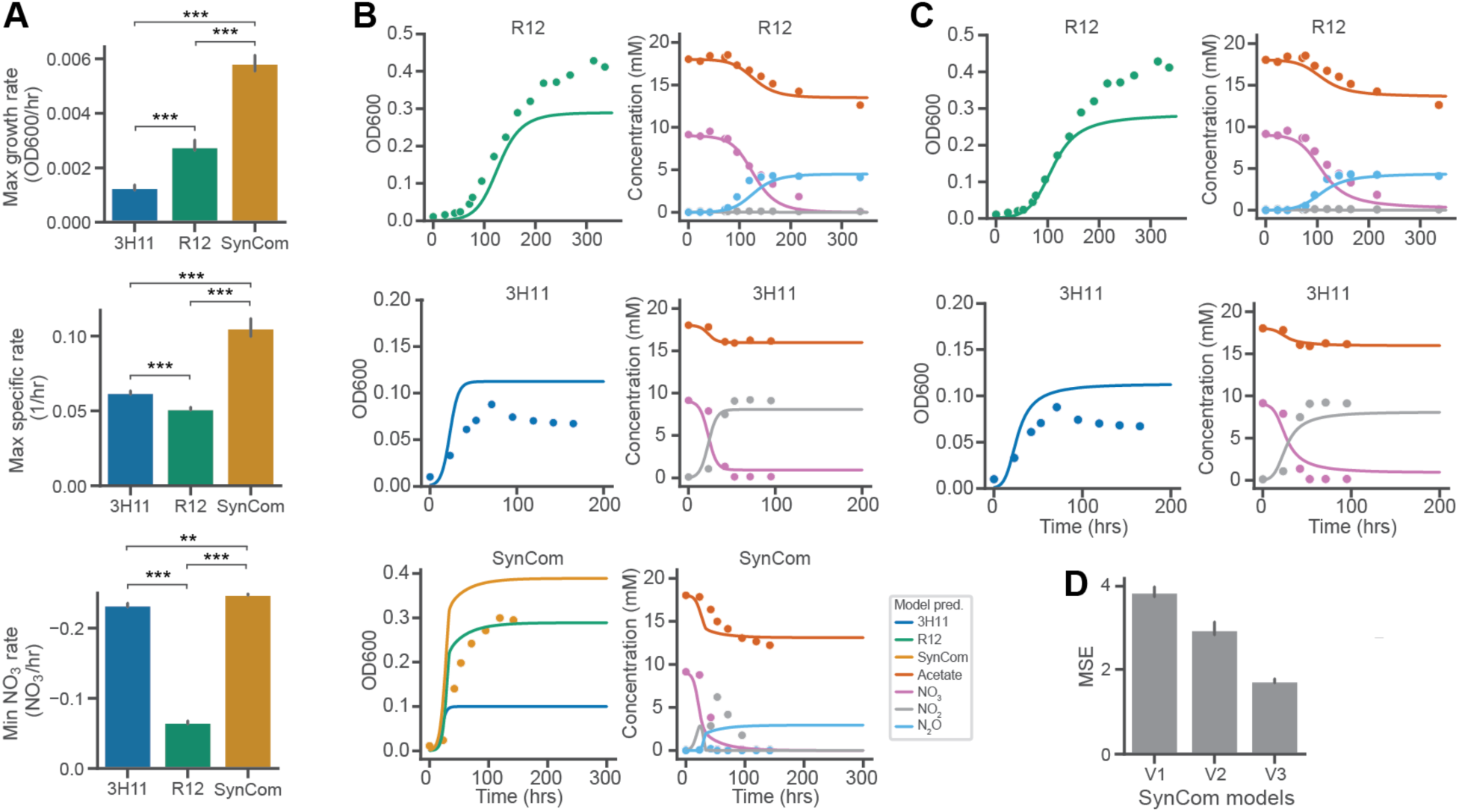
Kinetic modeling of individual monocultures and SynCom growth characteristics. (**A**) Comparison of monoculture and SynCom maximum growth rates, maximum specific rates and minimum NO_3_^−^ rate from the transcriptomics growth data. (**B**) Predicted growth dynamics and associated data for initial kinetic model formulations for R12, 3H11 and the SynCom. (**C**) Updated growth dynamics predictions with models which included NO_3_^−^ inhibition. Model predictions are represented using solid lines, data are averages across triplicate samples and are represented using circular points. SynCom models assume NO_2_^−^ and N_2_O are exchanged. Comparison of mean standard error (MSE) for three iterations of SynCom kinetic models. MSE computed across both growth and metabolite data. Bars indicate comparisons for which differences were significant using Welch’s t-test. *, *p*<0.05; **, *p*<0.01; ***, *p*<0.001.

**Supplemental figure 3.**
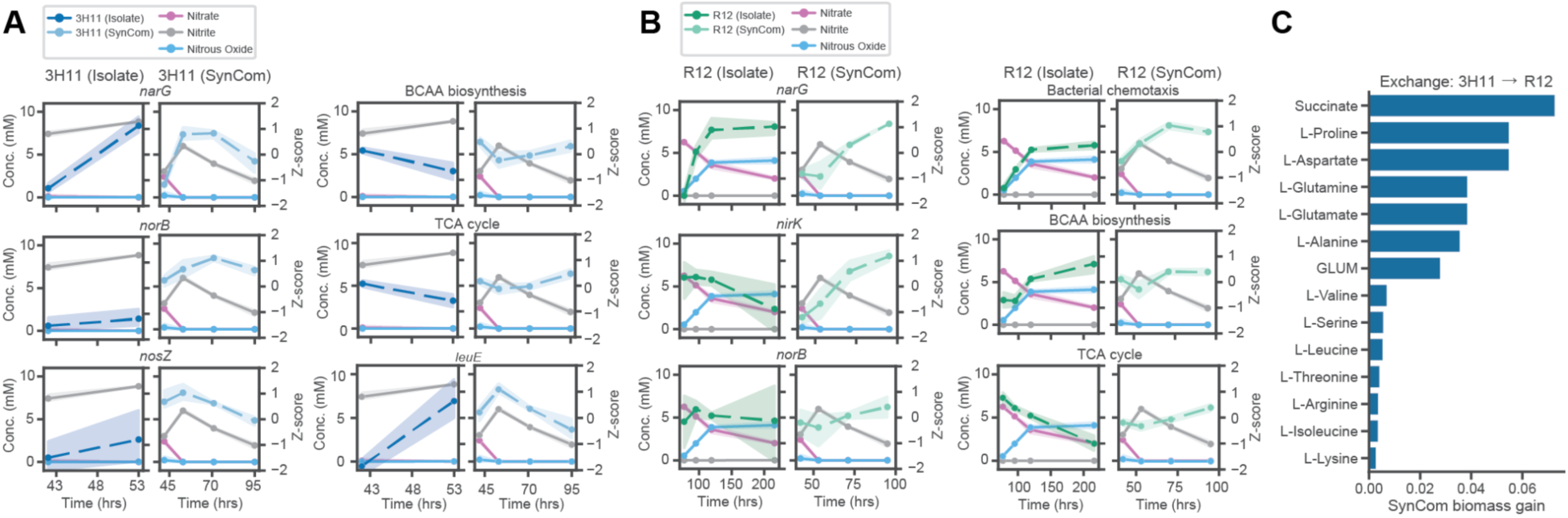
Time resolved expression dynamics of denitrification genes and select pathways. (**A**) 3H11 Expression dynamics for specific genes and pathways. (**B**) R12 Expression dynamics for specific genes and pathways. Points and trend lines represent averages of triplicate samples in the case of genes and averages across genes and samples in the case of pathways. Error bands display standard deviations. (**C**) Simulations with the constrains based metabolic model predict modest increase in biomass of the SyCom through exchange of amino acids and intermediates of central carbon metabolism.

**Supplemental figure 4.**
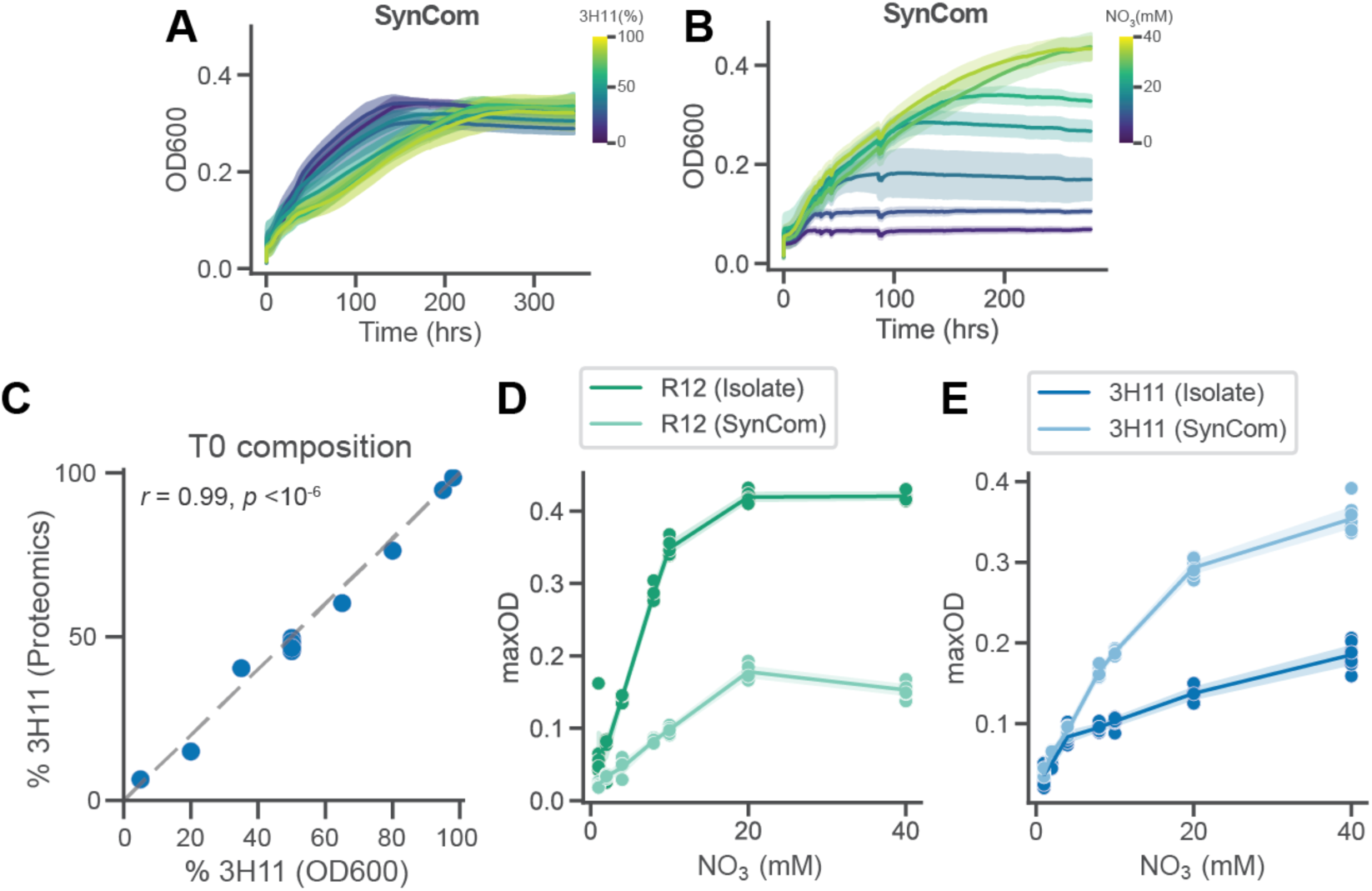
SynCom growth characteristics across variation in initial community composition and NO_3_ concentration. (**A**) Growth of the SynCpm across variation in initial community composition (3H11 %). Trendlines display average trajectories across eight replicates and error bands display standard deviations. Trendlines are colored by 3H11 %. (**B**) Growth of the SynCpm across variation in initial NO_3_ concentrations. Trendlines display average trajectories across eight replicates and error bands display standard deviations. Trendlines are colored by NO_3_ concentration. (**C**) Correlation between inoculum 3H11 % estimated by OD600 and proteomics. Dashed line indicates a 1:1 relationship. (**D**) Comparison of maximum OD600 achieved by R12 in monoculture vs R12 in the SynCom as a function of NO_3_ concentration. (**E**) Comparison of maximum OD600 achieved by 3H11 in monoculture vs 3H11 in the SynCom as a function of NO_3_^−^ concentration.

